# Protein unfolding thermodynamics predict multicomponent phase behavior

**DOI:** 10.1101/2023.05.26.542380

**Authors:** Neha Rana, Rukhillo Kodirov, Anisha Shakya, John T. King

## Abstract

An increasing number of proteins are known to undergo liquid-liquid phase separation (LLPS), with or without nucleic acids or partner proteins, forming dense liquid-like phases termed biomolecular condensates. This physical phenomenon has been implicated in the existence of cellular membraneless organelles as well as in the formation of pathological protein aggregates in several human diseases. While common structural features of proteins with a propensity to undergo LLPS have been well documented, currently there is no thermodynamic framework capable of predicting the phase behavior of native proteins. Here, we show that two fundamental thermodynamic properties associated with the unfolding of a native protein, change in heat capacity (*ΔC*_*unfold*_) and change in Gibbs free energy (*ΔG*_*unfold*_), are sufficient to predict the formation of multicomponent biomolecular condensates. We find that proteins with small *ΔC*_*unfold*_ and *ΔG*_*unfold*_ values, which indicate a native state that is thermodynamically similar to the fully unfolded state, promote LLPS. In contrast, proteins with large *ΔC*_*unfold*_ and *ΔG*_*unfold*_ values promote aggregation. We also demonstrate that the stability of the liquid-like condensate can be predicted from the proximity of a protein’s thermodynamic variable values to the phase boundary. This work elucidates a deep connection between single-protein thermodynamics and multicomponent phase behavior, and provides an avenue for predicting pathological droplet-aggregate transition.

**Significance Statement:** Biomolecular phase transitions, such as LLPS of proteins and nucleic acids, is emerging as an important concept in understanding the link between dysregulation of membraneless compartmentalization in cells and several human diseases. Currently, analysis of structural features such as intrinsic disorder and charged residue content is the “rule-of-thumb” to predict a protein’s propensity to undergo LLPS. There are several instances where this qualitative approach fails, potentially due to not accounting for a protein’s native structure. We demonstrate an empirical correlation between the unfolding thermodynamics of native proteins and their phase behavior, which enables a quantitative prediction of multicomponent LLPS. This novel approach, based on single-protein thermodynamics, has the potential to quantitatively predict pathological phase transitions of proteins in degenerative human diseases.

## Introduction

Cells can leverage selective, reversible assembly of proteins and nucleic acids via associative phase transition termed liquid-liquid phase separation (LLPS) as a means of spatiotemporal organization (1-3). This physical phenomenon is implicated in a variety of cellular processes, including chromatin organization (4-7), DNA repair (8-10), and cellular signaling (11, 12), with dysregulated phase transitions linked to various diseases (13-15). In general, assembly into stable liquid-like droplets is believed to be driven by non-specific interactions, including ion release, long-range electrostatic interactions, and short range cation-π and π−π interactions (16). With growing database of proteins shown to undergo LLPS, common structural features such as high content of charged residues, disordered low complexity prion-like domains (PLDs) (17), long intrinsically disordered regions (IDRs) at the N- or C-terminus (18, 19), and IDR linkers connecting structured regions in modular proteins (20, 21), all of which can promote multivalent interactions, have been consistently observed. These structural feature analyses have served as a general “rule-of-thumb” for predicting the phase behavior of proteins. To highlight the importance of IDRs in LLPS, many studies employ truncated proteins containing only the dominant IDR (17, 22, 23). The focus on IDRs has also led to a number of studies modelling LLPS based on long established theories of polymer solutions (18, 23, 24). Here, we demonstrate that such a qualitative rule-of-thumb that considers only certain structural features of a protein, is not sufficient to reliably predict LLPS of native proteins. In this work, we report a quantitative approach, based on protein unfolding thermodynamics, to predict multicomponent LLPS as well as the stability of the resulting condensates to external perturbations.

We investigate the multicomponent phase behavior of 13 native proteins with a range of structural features, including proteins with charged amino acid-rich long disordered terminal tails (histone proteins H1, H2A, H2A.Z1, H2B, and H3), RNA binding proteins (RBPs) with PLDs (TIA1 and TDP-43), intrinsically disordered proteins (prion protein (PrP) and Tau441), modular proteins (Nck1 and SH3 domain repeat of Nck2, (SH3)_4_), as well as almost completely structured proteins (lysozyme and glucose oxidase). Histone proteins, key architectural proteins in chromatin organization, have recently been shown to be involved in LLPS of chromatin and potentially contribute to the segregation of euchromatin and heterochromatin in the cell nucleus (4-7). TIA1 and TDP-43 localize in cellular biomolecular condensates such as stress granules and are implicated in RBP aggregation-related diseases (25, 26). Nck proteins, which are involved in signal transduction (27), have been shown to form signaling condensates at lipid membranes (28). Both PrP and Tau are extensively studied amyloid fibril forming proteins associated with neurodegenerative diseases (29, 30). The pathological aggregation of proteins like TIA1, TDP-43, PrP, and Tau, is thought to be mediated via LLPS (14, 25, 31, 32). To contrast with the phase behavior of IDR containing proteins, we also included the globular proteins lysozyme and glucose oxidase in our study.

Using a combination of differential scanning calorimetry (DSC) and in vitro phase separation assays, we find an empirical correlation between changes in heat capacity (*ΔC*_*unfold*_) and Gibbs free energy (*ΔG*_*unfold*_) upon unfolding and multicomponent phase behavior of the above-mentioned proteins in presence of oppositely charged homopolypeptides. Proteins with small *ΔC*_*unfold*_ and *ΔG*_*unfold*_ values promote LLPS, while proteins with large *ΔC*_*unfold*_ and *ΔG*_*unfold*_ values favor aggregation. *ΔC*_*unfold*_ quantifies the difference in local frequency of the free-energy minima associated to the native state of the protein and the unfolded state (33). As the configurational entropy of the unfolded state of the protein is maximum, *ΔC*_*unfold*_ is typically positive. *ΔG*_*unfold*_ quantifies the difference in free energy between the native and unfolded state (33) under standard conditions, and is likewise positive. The observed correlation indicates that proteins whose native structures are thermodynamically similar to their unfolded state promote the formation of stable liquid-like condensates, while proteins whose native structures are thermodynamically dissimilar to their unfolded state promote aggregation.

We also demonstrate that *ΔC*_*unfold*_ and *ΔG*_*unfold*_ values are sufficient to predict the stability of LLPS condensates to changes in temperature. We find that condensates formed by proteins that have *ΔC*_*unfold*_ and *ΔG*_*unfold*_ values close to the phase boundary are unstable to temperature changes. This can be attributed to the decreased solubility of the protein’s unfolded state with an exposed hydrophobic core (34), which promotes aggregation within the condensates (35). In contrast, proteins that have *ΔC*_*unfold*_ and *ΔG*_*unfold*_ values far from the phase boundary are stable to thermal perturbations, as unfolding does not lead to a significant change in solubility of the protein due to the small hydrophobic cores. These results show that we can utilize the basic thermodynamic parameters of a native protein to gain a predictive understanding of how proteins contribute to multicomponent LLPS and the stability of resulting condensates to external perturbation.

## Results and Discussion

### Empirical correlation between single-protein thermodynamics and phase behavior

To resolve the correlation between phase behavior and single-protein thermodynamics, we selected a series of proteins with structures ranging from nearly entirely disordered to entirely structured (**Fig. S1**). Phase separation assays were performed at pH = 7.4 and T = 22°C, unless stated otherwise. To maintain charge neutral condition between the polymers in the solution, either poly-L-aspartic acid (*n*=100, PLD100) or poly-L-arginine (*n*=100, PLR100) was added, depending on the net charge of the protein, such that the ratio of positively and negatively charged amino acids was 1 (N^+^/N^-^=1). We first compared multicomponent LLPS for all proteins used in the study as a function of salt concentration (**Fig. 1, S2, S3**). As expected, the LLPS of these proteins is dependent on the salt concentration. Most proteins are either in aggregate or solution phase at [NaCl] > 150 mM. At [NaCl]= 100 mM, 5 proteins (H1, H2A, PrP, (SH3)_4_, and Nck1) undergo LLPS to form condensate droplets in presence of oppositely charged polypeptides, while 8 proteins (H2A.Z1, H2B, H3, TIA1, TDP-43, Tau441, lysozyme, and glucose oxidase) either aggregate or remain in solution. At lower salt concentration, fewer proteins undergo LLPS.

**Fig. 1.**
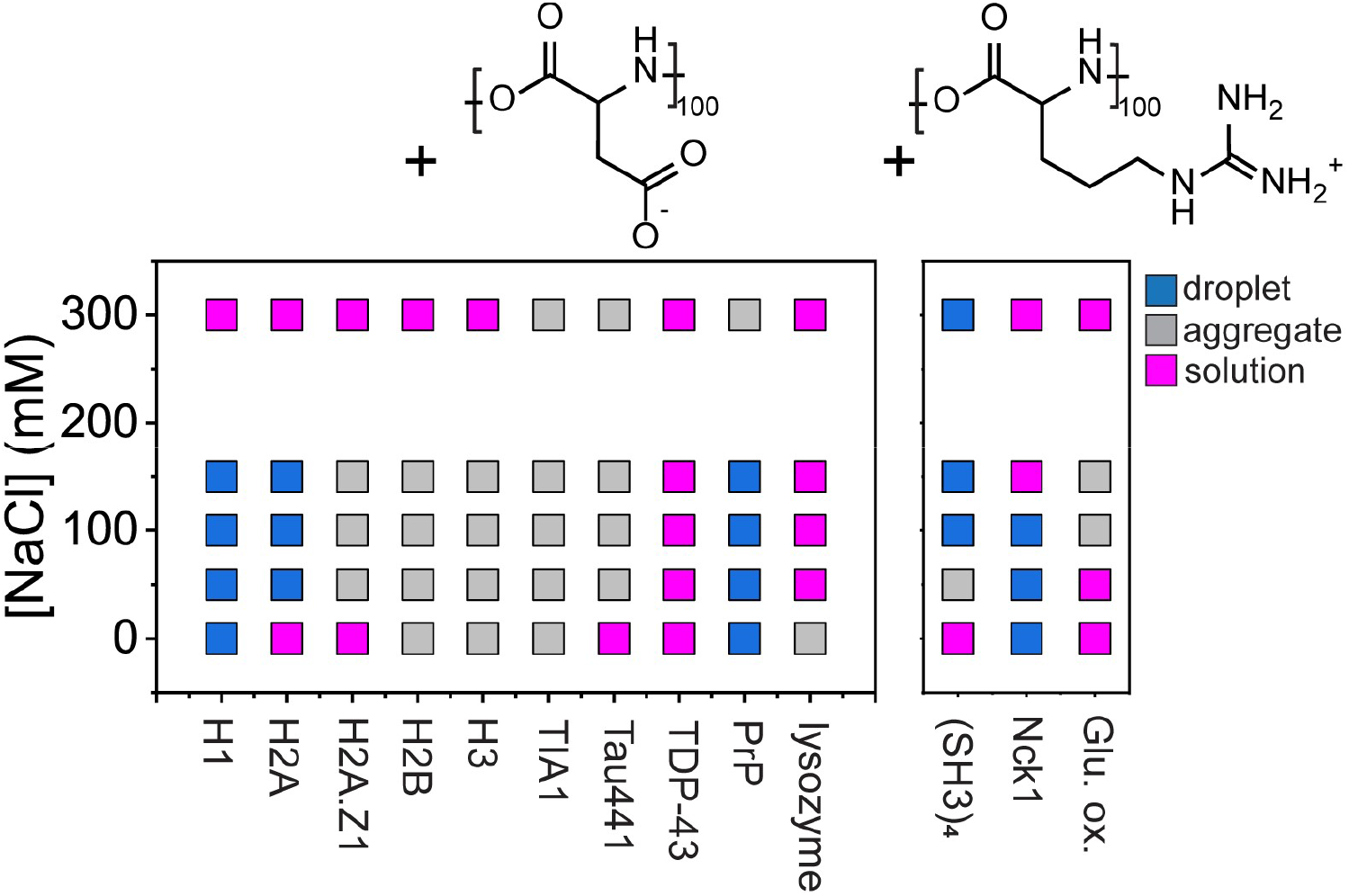
Multicomponent phase behavior of proteins in presence of oppositely charged polypeptides. **(left)** Proteins with excess positive charge mixed in solution with PLD100 and **(right)** proteins with excess negative charge are mixed in solution with PLR100 under charge neutral conditions (N^+^/N^-^ = 1). Phase behavior was measured as a function of [NaCl] ranging from 0 to 300 mM. Three distinct phases are observed: droplet (blue), aggregate (grey), or solution (pink) phase.

A scatter plot of phase behavior for these 13 proteins as a function of net charge and total number of disordered residues (**Table S1 and S2**) is shown in **Fig. 2a**. From the plot, it is evident that this qualitative rule of thumb is insufficient to predict multicomponent LLPS of proteins. Seeking an alternative approach, we use differential scanning calorimetry (DSC) to experimentally measure thermodynamic parameters associated with protein unfolding, namely *ΔC*_*unfold*_ and *ΔG*_*unfold*_, for each protein at pH = 7.4 and [NaCl] = 150 mM. **Fig. 2b** shows a scatter plot of protein phase behavior with *ΔC*_*unfold*_ and *ΔG*_*unfold*_ values. Here, we see a clear phase boundary: proteins with small *ΔC*_*unfold*_ and *ΔG*_*unfold*_ values promote LLPS, while proteins with large *ΔC*_*unfold*_ and *ΔG*_*unfold*_ promote aggregation.

**Fig. 2.**
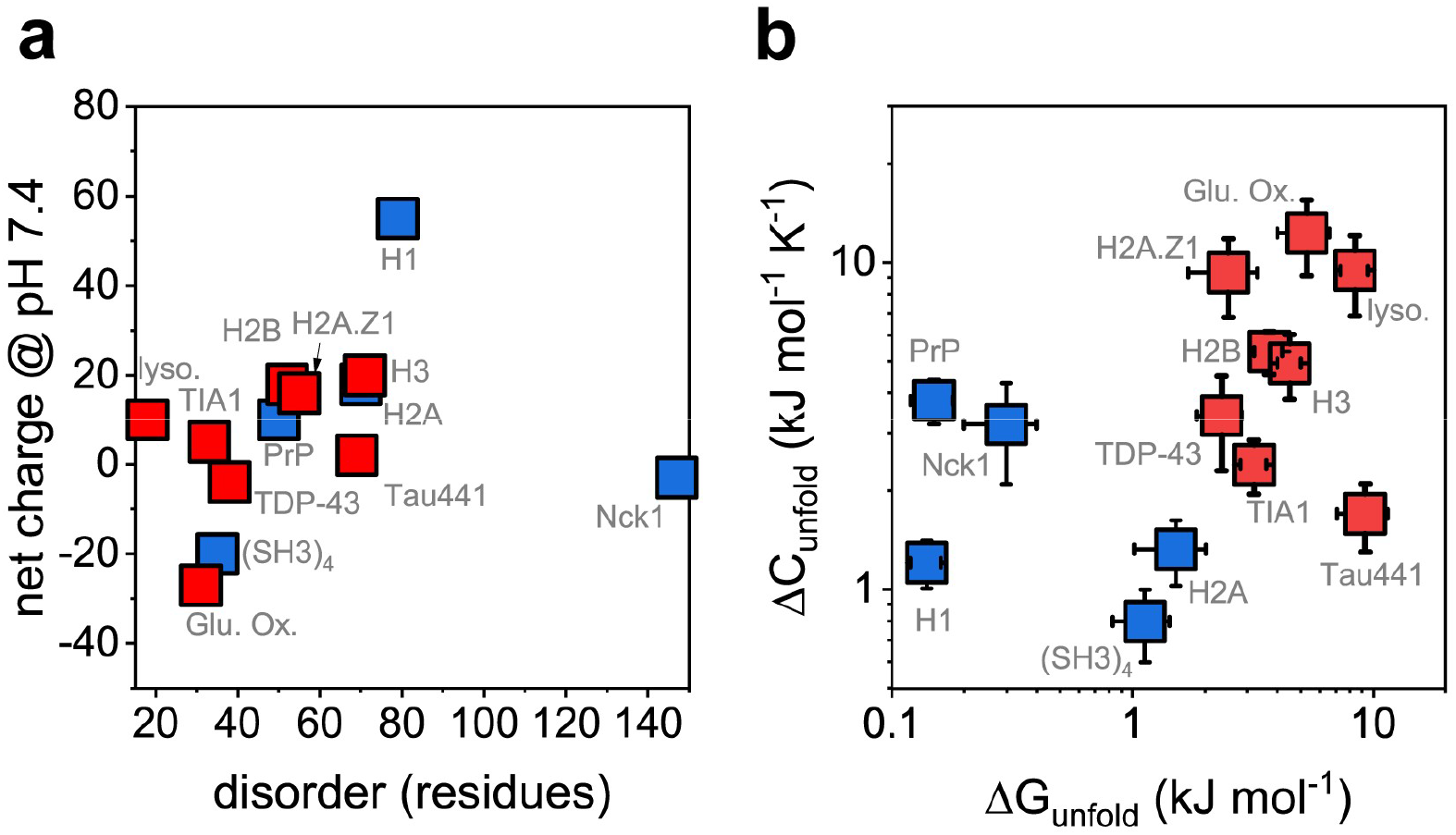
Empirical correlation between protein unfolding thermodynamics and phase behavior. **(a)** Scatter plot showing the phase behavior of proteins in the presence of oppositely charged polypeptides as a function of a protein’s net charge at pH 7.4 and number of intrinsically disordered residues. Proteins undergoing LLPS are shown in blue, while proteins that aggregate are shown in red. The phase behavior of a protein shows essentially no correlation with these structural feature parameters. **(b)** Scatter plot showing the phase behavior of proteins in the presence of oppositely charged polypeptides as a function of a protein’s unfolding thermodynamic parameters, *ΔC*_*unfold*_ and *ΔG*_*unfold*_. Here, there is a clear phase boundary is observed, where proteins with small *ΔC*_*unfold*_ and *ΔG*_*unfold*_ values promote LLPS, while proteins with large *ΔC*_*unfold*_ and *ΔG*_*unfold*_ values promote aggregation.

To rationalize the observed empirical correlation, we look at the physical meaning of these thermodynamic variables in reference to the underlying free energy surface of the protein. The *ΔG*_*unfold*_ of a protein represents the difference in thermodynamic potential between native and unfolded states. Assuming two-state model for unfolding, the *ΔG*_*unfold*_ can be determined from the melting temperature *T*_*m*_ and *ΔH*_*unfold*_ of a protein, both measured from DSC experiments (**Fig. S4**), using the Gibbs-Helmholtz equation,

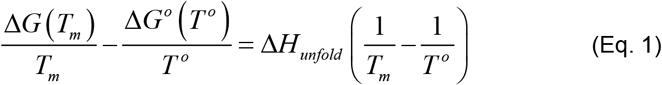

As *T*_*m*_ is defined as the temperature at which there are equal populations of proteins in the folded and unfolded states, the *ΔG* at *T*_*m*_ is 0. The *ΔG*_*unfold*_ is given by *ΔG*^*°*^ and can be determined from the simplified expression;

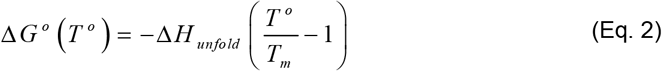

The constant-pressure heat capacity (*C*) of a protein has numerous mathematical definitions; the most relevant expression for this study relates *C* to the local curvature of the free energy basin of the protein (33),

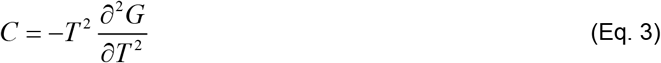

where *T* is the temperature. Therefore, *ΔC*_*unfold*_ represents the change in curvature of the free energy basin of the native structure relative to the fully unfolded protein. *ΔC*_*unfold*_ for a protein is nearly always positive, as the conformational freedom of the unfolded protein is always larger than that for the structured protein (36). Furthermore, protein unfolding is typically accompanied by the exposure of the hydrophobic core of the protein to hydration, which results in entropically unfavorable hydration and an increase in *C* (35). Structurally, proteins with small *ΔC*_*unfold*_ tend to have extended disordered regions, loosely folded domains, and/or folded domains with small hydrophobic cores (33, 36).

Taken together, *ΔC*_*unfold*_ and *ΔG*_*unfold*_ values characterize the thermodynamic similarity between the free energy surfaces of the native and unfolded states. Proteins with small *ΔC*_*unfold*_ and *ΔG*_*unfold*_ values are thermodynamically similar to the unfolded state, while proteins with large *ΔC*_*unfold*_ and *ΔG*_*unfold*_ values are thermodynamically dissimilar from the unfolded state. Thus, the scatter plot in **Fig. 2b** demonstrates that proteins with native state thermodynamic properties similar to the unfolded state promote LLPS, which is consistent with the general observation that intrinsically disordered domains drive phase separation.

To highlight the use of this thermodynamic analysis, we resolve a previously reported unexpected experimental observations on the phase separation of histone proteins with DNA (7, 37). The nucleosome core histones H2A, H2B, and H3 have similar size, degree of disorder, and excess net charge (**Table S2**). According to the general rule-of-thumb, we would anticipate these proteins to display similar phase behavior. However, H2A readily undergoes LLPS to form liquid-like droplets in presence of DNA (7) or oppositely charged polypeptides (**Fig. 1, S2**), while H2B, and H3 drives aggregation under the same conditions (**Fig. 1, S3**). We also tested H2A.Z1, a histone H2A variant with 60% sequence similarity to H2A (38), and find that it promotes aggregation (**Fig. 1, S3**). Our thermodynamic analysis of the proteins reveals that H2A has unexpectedly small *ΔC*_*unfold*_ and *ΔG*_*unfold*_ values compared to the H2B, H3, and H2A.Z1 (**Fig. 2b**). This analysis suggests that the folded domain of H2A is characterized by a less compact structure and smaller hydrophobic core than the structured domain of other core histones, making H2A more favorable to undergo LLPS.

We find that, of the proteins studied, the non-core histone, H1, has the lowest *ΔC*_*unfold*_ and *ΔG*_*unfold*_ values, consistent with a structure that has a small, folded core and extensive intrinsic disorder. Indeed, H1 strongly promotes LLPS in presence of negatively charged polypeptides (**Fig. S2**), as well as single-stranded and double-stranded DNA, and nucleosome arrays (3, 7, 37).

Two intrinsically disordered proteins, PrP and Tau441, implicated in degenerative diseases (29, 30), were also included in our thermodynamic analysis. Surprisingly, PrP, the smaller of the two proteins, is observed to have a larger *ΔC*_*unfold*_ value and smaller *ΔG*_*unfold*_ value than that of Tau441 (**Fig. 2b, Table S3**). This suggests that PrP can promote LLPS but is particularly susceptible to small fluctuations in the protein structure, and therefore protein solubility, which would destabilize the liquid-like condensate.

We also analyzed a set of proteins (TIA1, TDP-43, Nck1, (SH3)_4_) consisting of multiple folded domains bridged by disordered regions. Despite the modular nature of their structures, each protein shows a single melting temperature consistent with the two-state model of unfolding (**Table S3**). We find that while TIA1, TDP-43, and Nck1 have comparable *ΔC*_*unfold*_ values, Nck1 has a uniquely small *ΔG*_*unfold*_ value compared (**Fig. 2b**). As a result, we observe that Nck1 undergoes LLPS forming stable liquid-like droplets in presence of oppositely charged polypeptide (**Fig. 1, S2**), while TIA1 and TDP-43 aggregate (**Fig. 1, S3**). TIA1 has been previously shown to form droplets in presence of Zn^2+^ (39) or nucleic acids with specific binding sequences (40), while it forms fibrillar aggregates by itself at higher concentrations (26). Indeed, we find that in absence of any specific binding, TIA1 forms aggregates, both by itself and in presence of PLD100. Similarly, TDP-43 has been shown to undergo LLPS in presence of specific binding partners (41, 42), Here, we find that it preferentially aggregates in presence of non-specific interactions. In the case of (SH3)_4_, we observe a uniquely small *ΔC*_*unfold*_ value and hence is observed to undergo LLPS forming liquid-like droplets (**Fig. 1, S2**).

For completeness, we also analyzed two highly structured proteins, lysozyme from hen egg white and glucose oxidase (**Fig. S1, S3**). As expected from the structure of the proteins, both have high *ΔC*_*unfold*_ and *ΔG*_*unfold*_ values (**Fig. 2b**). The observation that the proteins do not undergo multicomponent LLPS in presence of oppositely charged polypeptides at low salt concentrations is therefore expected.

### Reversibility of protein-polypeptide condensates

In the previous section, we demonstrated the thermodynamic criteria by which native proteins undergo LLPS in the presence of polypeptides. In this section, we show how a protein’s thermodynamic parameters impact the stability (*i*.*e*., reversibility) of the resulting multicomponent condensates.

We first studied the response to heating/cooling of the condensates of three proteins whose *ΔC*_*unfold*_ and *ΔG*_*unfold*_ values fall well-below (H1), well-above (H3), and close to the phase boundary (H2A) (**Fig. 2b**). As measured using DSC, the *T*_*m*_ of H1, H2A, and H3 are 39°C, 42°C, and 66°C, respectively (**Table S3**). **Fig. 3a, top panel left to right**, shows droplets (formed after mixing H1 and PLD100 solutions incubated at 20°C) heated gradually to 44°C. **Fig. 3a, bottom panel right to left**, shows droplets (formed after mixing H1 and PLD100 solutions incubated at 44°C) cooled gradually to 20°C. Following incubation at 20°C, H1-PLD100 mixtures form smooth and circular droplets indicating a liquid-like character. As the sample is heated above the *T*_*m*_ of the protein, the droplets remain stable (*i*.*e*., liquid-like), with the surface coverage increasing slightly at elevated temperatures (**Fig. 3b**). When the components are incubated at 44°C (above the *T*_*m*_ of H1), liquid-like droplets are still formed, and the surface coverage decreases as the system is cooled to 20°C (**Fig. 3b**), indicating the reversible nature of the assembly.

**Fig. 3.**
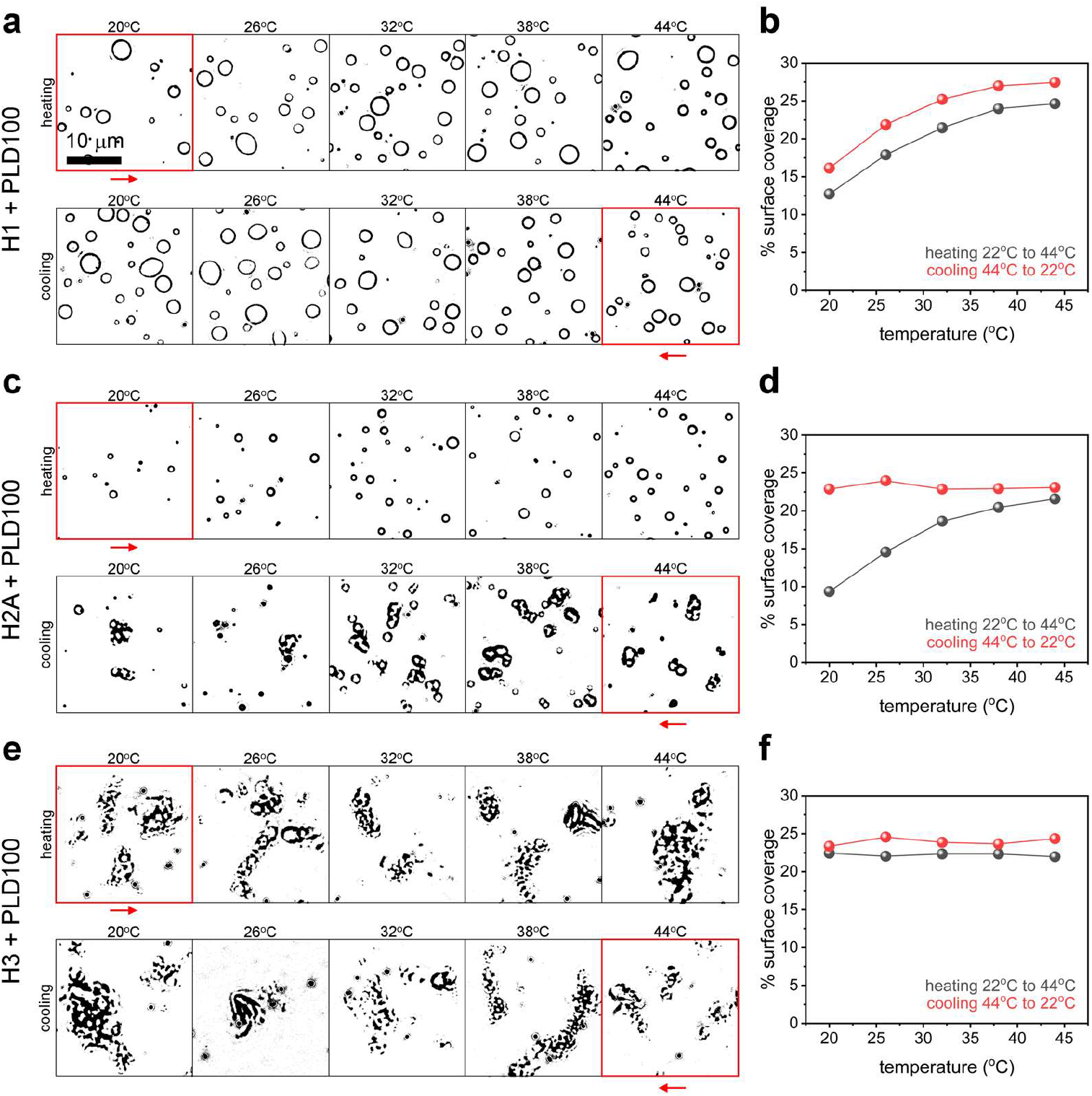
Irreversible aggregation in protein-polypeptide condensates. (**a**) Temperature dependent phase separation of H1-PLD100 mixtures and (**b**) corresponding surface coverage statistics. H1 shows reversible LLPS in presence of polypeptide when incubated both below and above *T*_*m*_. (**c**) Temperature-dependent phase separation of H2A-PLD100 mixtures and (**b**) corresponding surface coverage statistics. H2A forms stable liquid-like droplets when the protein is incubated below *T*_*m*_, but drives irreversible aggregation when incubated above *T*_*m*_. (**e**) Temperature-dependent phase separation of H3-PLD100 mixtures and (**b**) corresponding surface coverage statistics. H3 drives irreversible aggregation both below and above its *T*_*m*_, and therefore shows no temperature dependence.

**Fig. 3c** shows the same assay for H2A-PLD100 condensates. Unlike H1, the *ΔC*_*unfold*_ and *ΔG*_*unfold*_ values of H2A sit near the phase boundary (**Fig. 2b**). We observe liquid-like droplet formation only when the components of the mixture were incubated at 20°C, which is below the *T*_*m*_ of H2A (**Table S3**). When incubated above the *T*_*m*_ of H2A, the mixture forms irreversible aggregates that do not transition to liquid-like droplets upon cooling (**Fig. 3d**). In the case of H3, the *ΔC*_*unfold*_ and *ΔG*_*unfold*_ values lie above the phase boundary (**Fig. 2b**). Therefore, H3 forms aggregates with PLD100 when mixed at both 20°C and 44°C (**Fig. 3e**), with the aggregates being irreversible with respect to temperature changes (**Fig. 3f**).

The reversibility of condensate formation is critical to its function and stability (23). The relative stability of protein-polypeptide condensates for the proteins H1 and H2A can be understood from the protein’s *ΔC*_*unfold*_ value and its proximity to the phase boundary (**Fig. 2b**). The exposure of the hydrophobic core to hydration water upon denaturation is a major contributor to the positive *ΔC*_*unfold*_ values observed for protein unfolding (33, 36), as well as the driving force for hydrophobic assembly of the significantly less soluble unfolded protein (34). While both H1 and H2A have *ΔC*_*unfold*_ values compatible with LLPS, we find that the *ΔC*_*unfold*_ value of H1 is significantly smaller than that of H2A (**Fig. 2b**). The exceptionally small *ΔC*_*unfold*_ value of H1 results in the protein remaining highly soluble at all temperatures, even above the *T*_*m*_. In contrast, the *ΔC*_*unfold*_ value of H2A is near the phase boundary and is therefore capable of driving aggregation in the unfolded state.

### Thermal stability of protein-polypeptide condensates

In this section, we show that the temperature at which condensates become unstable and undergo irreversible aggregation is predicted from the protein’s *ΔC*_*unfold*_ value. The *ΔC*_*unfold*_ of a protein depends on both the interactions between amino acids in the protein in its native state and the hydration of the hydrophobic core upon unfolding (33). As the exposure of a protein’s hydrophobic core is accompanied by a decrease in solubility (34), the *ΔC*_*unfold*_value should reliably reflect a protein’s tendency to irreversibly aggregate when unfolded.

**Fig. 4a** shows the multicomponent phase behavior of 5 droplet forming proteins (PrP, Nck1, H2A, H1, and (SH3)_4_) as a function of mixing temperature, ranging from 20°C to 44°C. PrP, whose *ΔC*_*unfold*_value lies near the phase boundary and is the largest value amongst the 5 proteins (**Fig. 2b**), transitions from forming stable liquid-like droplets to irreversible aggregates at 32°C. As the *ΔC*_*unfold*_ value of the proteins decreases, we find the droplet-aggregate transition temperature occurs at increasingly high temperatures (**Fig. 4a**). This trend culminates with H1 and (SH3)_4_, which have exceptionally small *ΔC*_*unfold*_ values and hence continue to form stable liquid-like droplets at all temperatures studied.

**Fig. 4.**
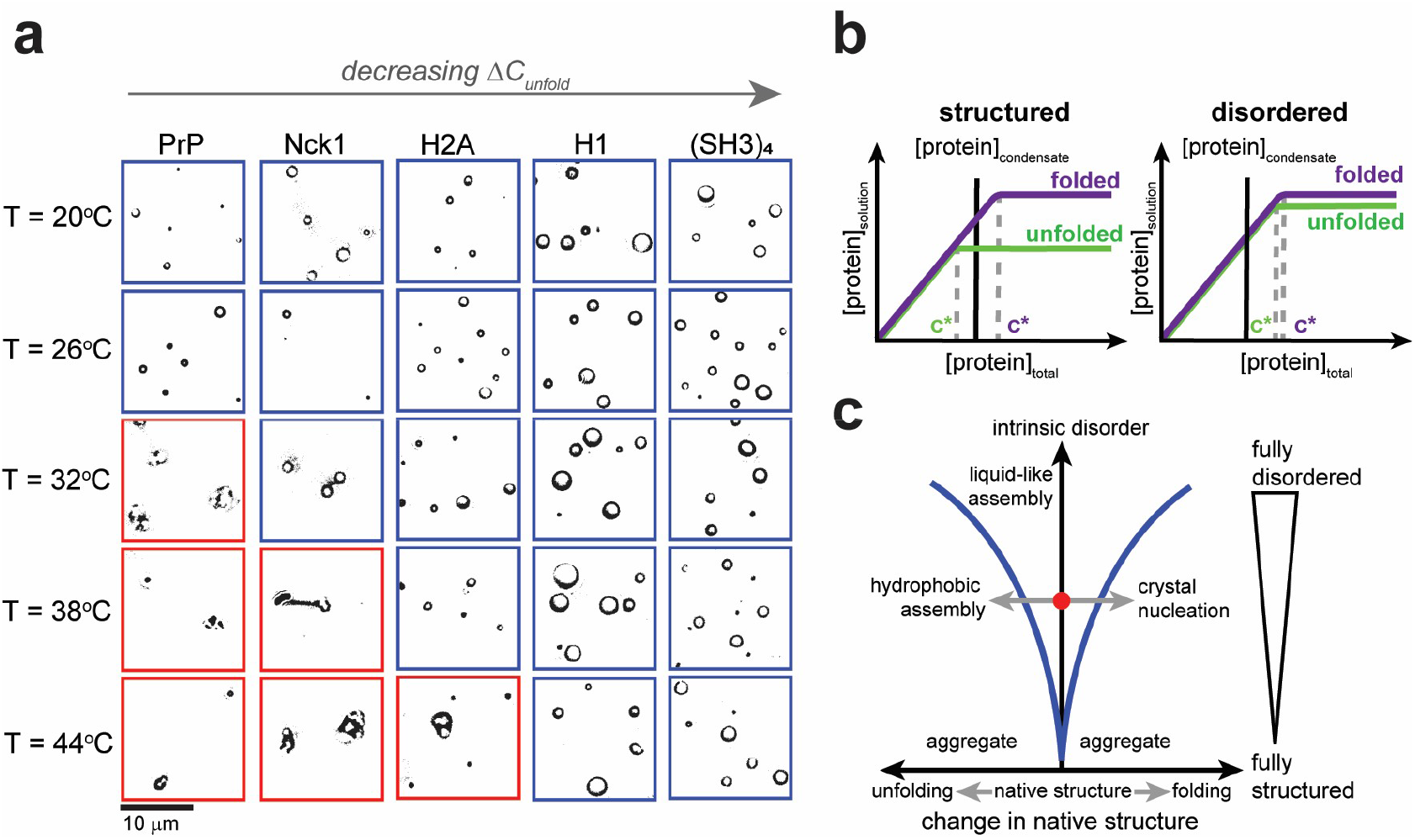
Thermal stability of protein-polypeptide condensates as a function of *ΔC_unfold_*. **(a)** Proteins with large *ΔC*_*unfold*_ values undergo irreversible aggregation at lower temperatures compared to proteins with small *ΔC*_*unfold*_ values. The results demonstrate that a protein’s heat capacity is the key parameter determining thermal stability of condensates. **(b)** Effect of protein unfolding on the solubility saturation concentration (*c\**). For native proteins, the concentration within the condensate is expected to be below the *c\**. Upon unfolding, proteins with extensive folded domains undergo a significant change in solubility as the hydrophobic core is exposed, leading to a *c\** that is potentially below the protein concentration within the condensate, thus driving aggregation. In contrast, proteins that contain only small folded domains undergo small changes to *c\** upon unfolding, and therefore the droplets remain stable even above the *T*_*m*_ of the protein. **(c)** Effect of protein structure deviations from the native state on droplet stability. For structured proteins, minor changes to the proteins structure promote aggregation, either due to partial unfolding or due to increased structure. For disordered proteins, the native state promotes formation of liquid-like condensates, which remain stable for even relatively large changes to the protein structure away from the native conformation.

Taken together, these results suggest that the solubility difference between a scaffold protein’s native state and the unfolded state determines the thermal stability of a biomolecular condensate (**Fig. 4b**). It is generally thought that LLPS condenses proteins to concentrations that approach the saturation concentration of the protein (*c\**). While the high concentrations of proteins potentially facilitate the function of the condensate (43), it can also drive unwanted aggregation if the protein partially or fully unfolds. For proteins with extensive folded domains, this structural change leads to a significant decrease in *c\**. When *c\** decreases below the condensate concentration it drives aggregation. In contrast, largely unstructured proteins experience little change in *c\** upon unfolding and can therefore remain in stable liquid-like droplets well above their *T*_*m*_.

The dependence of droplet stability on deviations from the native structure of the protein is illustrated in **Fig. 4c**. Proteins with extensive intrinsic disorder have the largest configurational space for LLPS driven droplet formation. As these proteins lack large hydrophobic cores, even full unfolding does not significantly destabilize the droplet. In contrast, highly structured proteins have a narrow configurational space for LLPS that is centered around the native state. Therefore, even small changes to the protein’s structure can induce instability through changes in the solubility of the protein (**Fig. 4b, left**). Our results demonstrate that our approach, which utilizes thermodynamic analysis of protein unfolding, provides a clear picture of droplet stability for various proteins having a range of structural features. This approach allows quantitative prediction of the response of a biomolecular condensate to changes in its environmental conditions due to various factors that affect the protein structure.

## Conclusions

The droplet to aggregate transition of biomolecular condensates is thought to be associated with several degenerative diseases. A quantitative understanding of the factors that drive stability of the liquid-like state of biomolecular condensates enables a predictive understanding of the pathological phase transitions of proteins implicated in such diseases. In this work, we seek to provide a general framework for predicting phase behavior and stability of biomolecular condensates from a thermodynamic characterization of the native protein. We report an empirical correlation between the *ΔC*_*unfold*_ and *ΔG*_*unfold*_ values of a protein and its propensity to promote LLPS over aggregation in presence of oppositely charged polypeptides. We rationalize this correlation by considering the free-energy profile of protein folding, where *ΔC*_*unfold*_ and *ΔG*_*unfold*_ values correspond the change in local curvature and depth of the free-energy basins of the folded and unfolded state. Proteins with small *ΔC*_*unfold*_ and *ΔG*_*unfold*_ values that favorably undergo LLPS are thermodynamically similar to their unfolded state. This result provides an underlying quantitative framework for understanding the general observation that proteins found in LLPS condensates are typically rich in disordered regions. Furthermore, we find that the proximity of the *ΔC*_*unfold*_ value to the phase boundary is a reliable predictor of the stability of biomolecular condensates upon changes to temperature. Proteins with *ΔC*_*unfold*_ value near the phase boundary typically have significant hydrophobic core to the structured region of the protein, which drives hydrophobic aggregation upon exposure to the solvent. Our results provide a quantitative framework for understanding multicomponent LLPS of different classes of proteins based on their unfolding thermodynamics. This framework can be generally applied to understanding the droplet-aggregate transition of a biomolecular condensate when subjected to any perturbations that affects its scaffold protein’s structure.

## Acknowledgements

We thank Malvern Analytics for assistance with DSC measurements. We thank the Schärer Lab for help with protein purification. We also thank the taxpayers who supported this work through the Korean Institute for Basic Science, project code IBS-R020-D1.

## Supporting Information

**Fig. S1.**
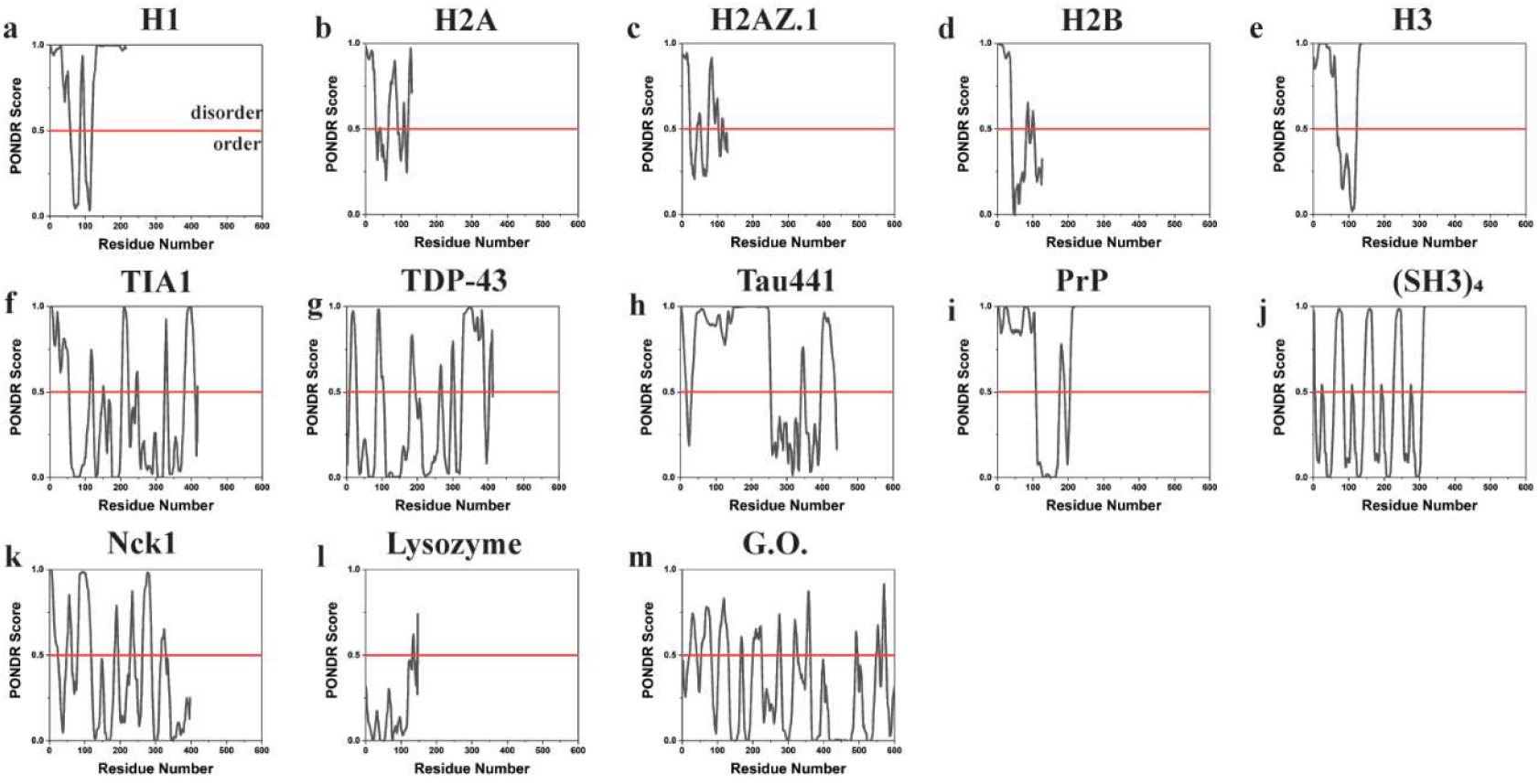
Prediction of disordered content of proteins using Predictor of Natural Disordered Regions (PONDR). The x-axis represents the total number of residues in each protein. The y-axis represents the PONDR score. A score greater than 0.5 indicates significant structural disorder.

**Fig. S2.**
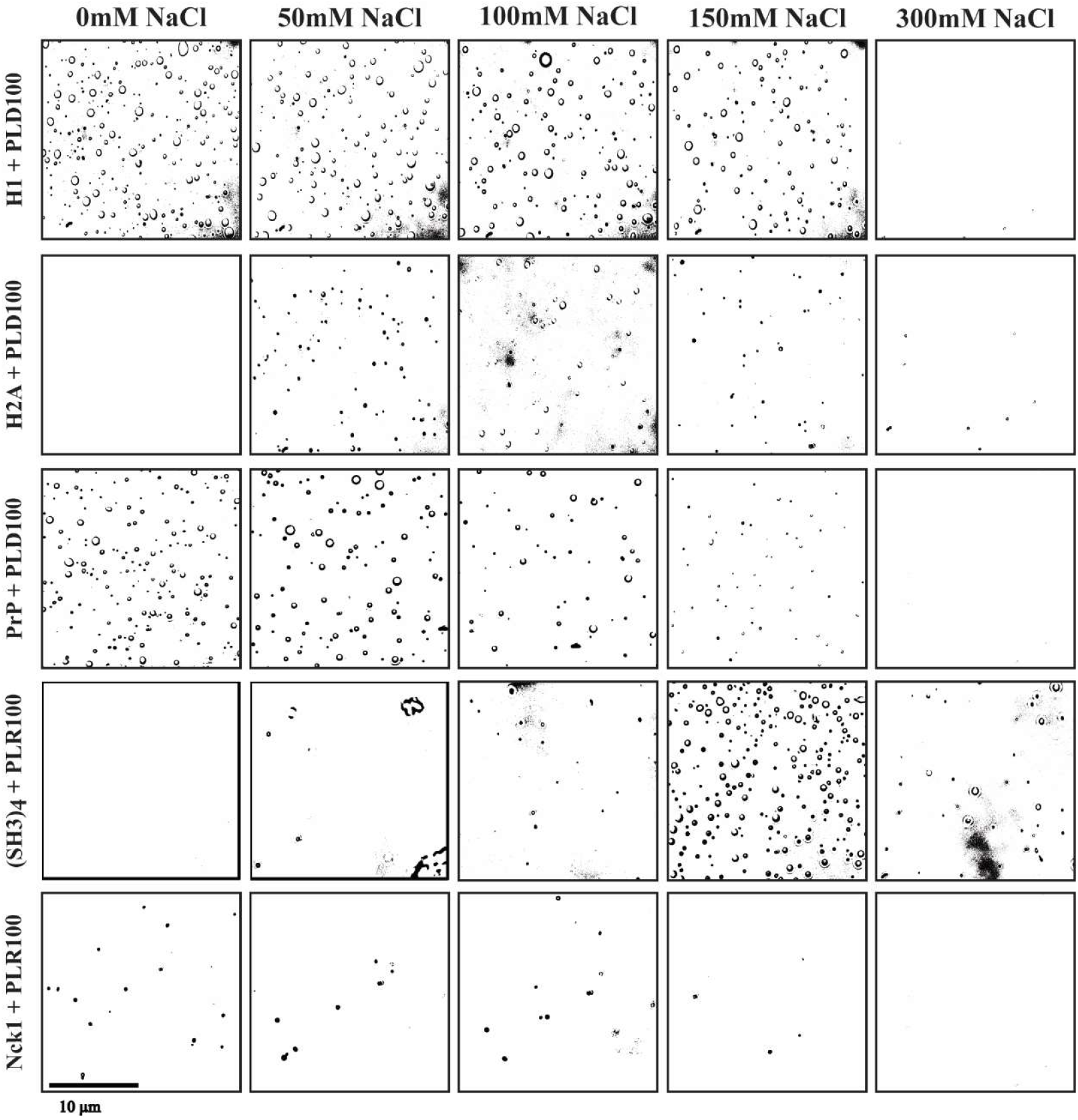
Phase separation assay at different salt concentrations for 5 droplet-forming proteins (H1, H2A, PrP, (SH3)_4_, Nck1) in presence of charged polypeptides PLR or PLD.

**Fig. S3.**
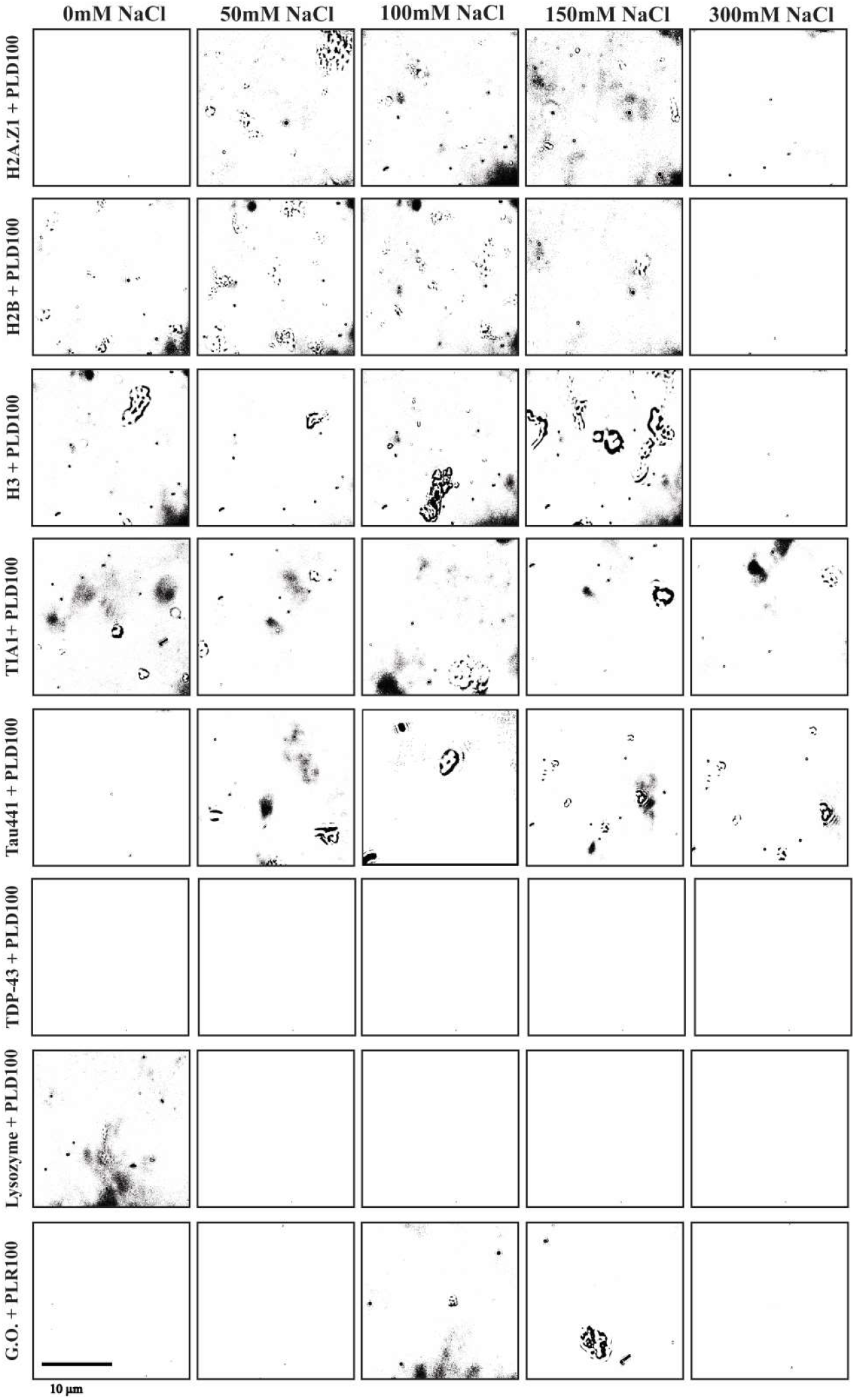
Phase separation assay at different salt concentrations for 8 aggregate-forming proteins (H1, H2A, PrP, (SH3)_4_, Nck1) in presence of charged polypeptides PLR or PLD.

**Fig. S4.**
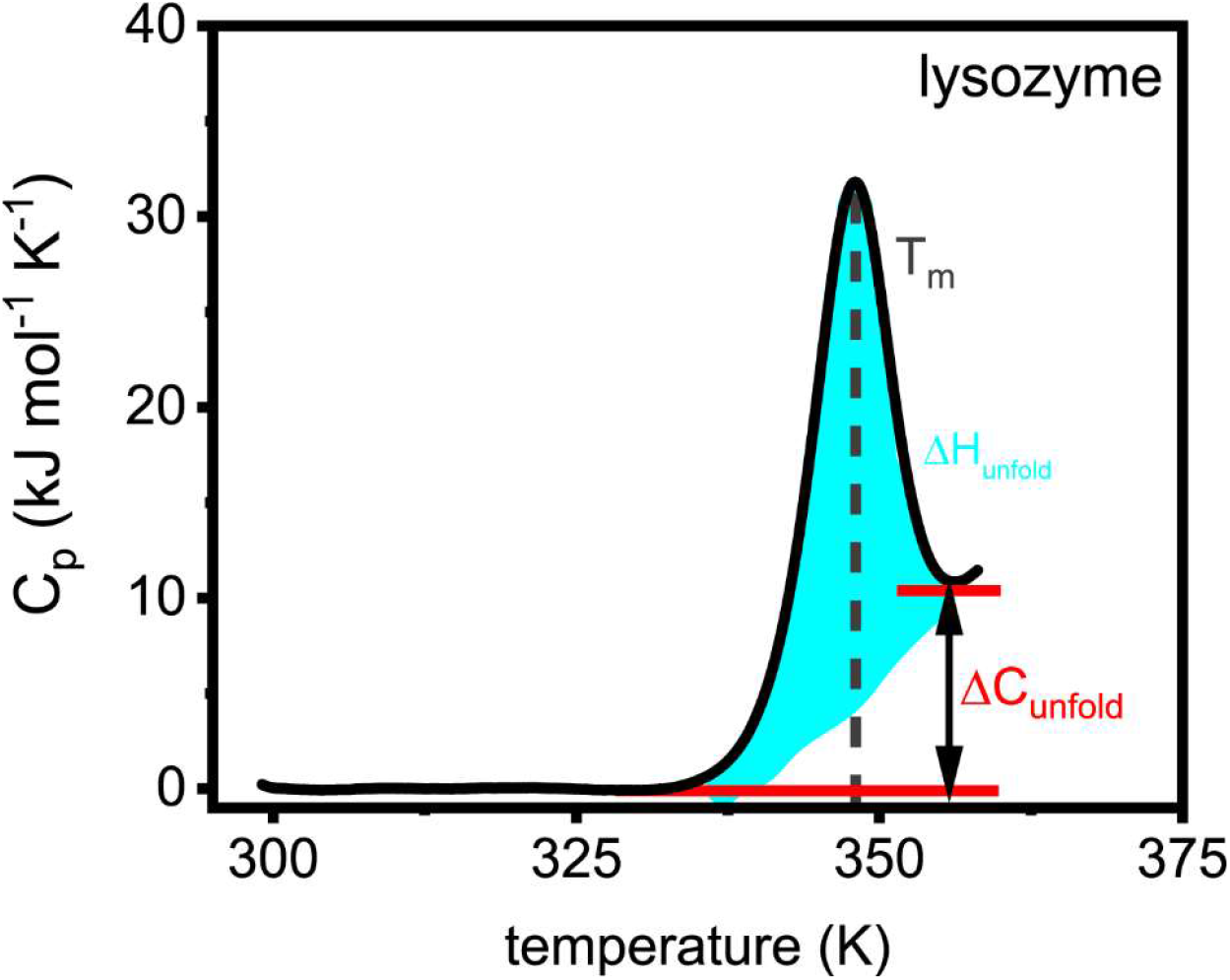
Differential scanning calorimetry (DSC) data for lysozyme. The melting temperature *T*_*m*_, *ΔH*_*unfold*_, and *ΔC*_*unfold*_ are directly extracted from the DSC data. The *ΔG*_*unfold*_ value is calculated using the *ΔH*_*unfold*_ and the Gibbs-Helmholtz equation.

